# Identification and classification of all Cytochrome P450 deposits in the Protein Data Bank

**DOI:** 10.64898/2026.03.17.712328

**Authors:** Piotr Śmieja, Marta Zadrożna, Khajamohiddin Syed, David Nelson, Dominik Gront

## Abstract

Cytochrome P450 monooxygenases (CYPs/P450s) form a highly diverse enzyme superfamily central to biotechnology, pharmacology, and environmental science. Despite the large number of available structures, identifying and comparing P450 entries in structural repositories remains challenging due to their extreme sequence divergence and inconsistent annotation practices. In particular, many deposits lack the standardized nomenclature (*CYPid*) and rather rely on legacy or author-defined *common names* (like *P450cam*, *P450BM-3* and *P450-PCN1*), which are often inconsistent in formatting and specificity. This is particularly difficult for a superfamily as sequentially diverse as P450s. This hinders reliable retrieval and cross-referencing, making even identification all P450 structures in the database nontrivial. To overcome these obstacles, we developed a structure-guided discovery and validation workflow combining keyword search, Hidden Markov Models, and structural alignment, enabling robust detection and annotation. This strategy identified 1,513 deposits representing 674 unique sequences. All sequences were reannotated using the P450Atlas server and manually verified, confirming high assignment accuracy. In the process, we have also identified five new CYP subfamilies. The resulting dataset constitutes the first rigorously curated, structure-linked registry of P450 enzymes, integrated into a publicly accessible resource and supported by an automated pipeline that periodically scans newly released entries. By unifying structurally validated identification with standardized CYP nomenclature, this work establishes a reliable framework for accurate retrieval, comparison, and future large-scale analyses of P450 enzymes.

## 1 Introduction

### 1.1 Cytochrome P450s

Cytochrome P450 monooxygenases (CYPs or P450s) are heme-containing enzymes that are ubiquitously distributed across all domains of life [1]. The term cytochrome *P450 monooxygenases* reflects their defining properties: cytochrome denotes a heme-bound protein, *P450* refers to the characteristic absorption peak at 450nm exhibited by the reduced enzyme when bound to carbon monoxide, and *monooxygenase* describes their canonical catalytic activity, the incorporation of one oxygen atom into a substrate [2, 3, 4, 5]. A detailed account of the historical discovery and nomenclature of these enzymes is available elsewhere [6]. Despite the name *monooxygenases*, P450 enzymes are capable of catalyzing a wide range of chemically diverse reactions, often with high regio- and stereospecificity, while also displaying notable substrate promiscuity. One of the earliest isolated members of this family was cytochrome P450cam (CYP101A1) from *Pseudomonas putida*, which catalyzes camphor hydroxylation and is therefore also known as camphor hydroxylase [7, 8]. CYP101A1 subsequently became a model P450 system and was the first P450 enzyme to be crystallized, providing key structural insights into the biochemical and mechanistic properties of this enzyme superfamily [9]. Since their initial discovery, a vast number of P450 enzymes have been identified across diverse organisms [10].

### 1.2 CYPid classification

In 1987, in response to the growing interest in the P450 superfamily, the first cytochrome nomenclature system was proposed, where a given gene name would consist of the *P450* prefix, a Roman numeral denoting a family, a capital letter denoting a subfamily, and an Arabic number for a specific gene [1]. At that time, however, the entire list of P450s consisted of just 65 entries. In 1989, the follow-up article was published, introducing the updated nomenclature, as well as new family and subfamily definitions [11]. The new system replaced the *P450* prefix with *CYP* (or *Cyp*) and required the family to be denoted by an Arabic number instead of a Roman numeral. In the following years, the updated lists of cytochrome families were published, with no significant changes to the naming scheme [12, 13, 14]. In 1994, when the space of known P450 sequences had become too large to conveniently publish in the traditional format, a website hosting all families and subfamilies was released (the first article to reference it was published in 2009 [15]). The nomenclature outlined above and illustrated in Figure 1 remains the current standard for naming cytochrome P450 sequences. In this work, identifiers formatted according to this convention are referred to as *CYPid*. A P450 family is typically defined based on sequence identity of more than 40% between each pair of member amino acid sequences, while the cut-off for subfamilies is 55%. These categories, however, are meant to reflect the evolutionary relations, and the individual decisions concerning particular genes may violate these rules [16].

**Figure 1:**
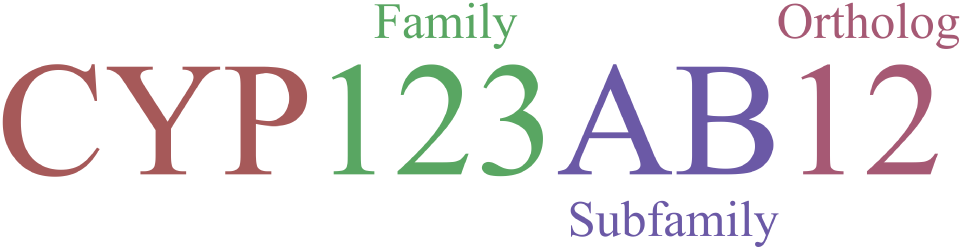
Breakdown of the CYPid cytochrome classification category

Traditionally, the family and subfamily for a newly identified P450 sequence had to be determined manually by a member of P450 nomenclature committee, creating a bottleneck in research. To answer this issue, a subfamily assignment server *P450atlas* [10] was published, providing a new, fast, and automated method for identifying and classifying P450 sequences. It uses Hidden Markov Model profile search to identify the best-fitting family and subfamily, followed by sequence alignment with the representative proteins for each candidate family’s ortholog groups. As a result, user obtains a list of sorted possible matches with values of the sequence identity between the query and the representative sequences.

### 1.3 Non-standard nomenclature

Many cytochromes have specific names, either besides, or instead of, CYPid. In some cases, these names predate the adoption of CYPid, as well as the family and subfamily classification. They are usually derived from their origin (e.g. *P-450BM3*) or enzymatic function (e.g. *P450CAM*), rather than the sequential similarity and evolutionary relations. In this article we refer to such names as *common names* or *aliases*. Despite their popularity, there are important downsides of such names. While they typically follow patterns like *P450-XXX*, *P450xxx*, *P450 XXX* or *P450(XXX)*, where *XXX* can denote a substrate, host organism or function, there is no well-established standard. Sometimes the suffix *XXX* is in the subscript (*P450_xxx_*) and *P450* may be hyphenated (*P-450*). Often the same name is typed in several different ways in various articles e.g. *P450CAM* in one publication and *P450(cam)* in another. Moreover, such names can be ambiguous: a single name can refer to multiple different P450s and a single P450 can have more than one name. We demonstrate this in section 3.3; for example CYP3A4 is referred to both as *P450-PCN1*, *HLp* and *NF-25*, and additionally *P450-PCN1* can also mean CYP3A1.

### 1.4 Research motivation

Since the first P450 crystallized in 1985 [9], structures of many P450 enzymes belonging to different families and subfamilies have been determined, and deposited in the Protein Data Bank (PDB)[17, 18]. However, these deposits are not consistently and sufficiently labelled according to the P450 classification and nomenclature system. In some cases no subfamily is listed, sometimes the provided subfamily is incorrect and in multiple cases no family is given at all. Often the *CYPid* format is not followed and instead the family is listed as *P450 xxx* (e.g. *P450 BM3*). As mentioned, many common names are in use, following many different formats, often instead of CYPid.

Such nomenclature was justified in the 1980s, with the limited number of known P450s, and the CYPid classification not yet established. Currently, however, with over 164,000 known P450 sequences and over 1,500 structures, this approach strongly inhibits research. These issues negatively impact searchability and reduce the reliability of any automated literature-mining procedures that depend on consistent nomenclature. (e.g. standard keyword search in RCSB cannot find all P450 deposits, while returning numerous false positives). Meanwhile, the superfamily’s diversity makes the commonly available sequence-based methods like BLAST insufficient and impractical.

Therefore, precise, standardized and unambiguous annotations in form of CYPid are highly needed. Yet, even now, as shown by the data presented in this publication, structures and sequences are often deposited, accompanied solely by a common name, rather than the proper CYPid. All this leads to misrepresentation of the P450s by researchers. This study aims to address this research gap by finding and analyzing all P450 structures in the PDB and providing the correct CYPid for each deposit in a publicly accessible and up-to-date database. Simultaneously, the project enhances the P450atlas sequence database with structural information, while also demonstrating its accuracy and usefulness.

## 2 Materials and methods

### 2.1 Tools

For file preparation and conversion, including chain extraction, we used a combination of BioShell 3.0 [19] and PyMol version 3.1.0 [20] packages. For clustering and sequence alignment we used programs of the BioShell package. Structural alignments were performed using the TMalign [21] program. Analysis was done using Python and the images were prepared using Matplotlib [21] version 3.10.0, Blender version 4.3.2 [22], and Visualife [23] packages.

### 2.2 Dataset and file preparation

For each deposit in the RCSB PDB (accessed on 28.07.2024) we obtained the corresponding file in the mmCIF format, extracted all the polymer entities and, for each one, we chose the longest (most complete) chain available. Typically, in each deposit, there would only be one entity with a cytochrome. In rare cases, in which there were more P450-containing entities, the enzymes were almost the same with minor differences, such as a his-tag, or a few missing termini residues. In all cases, we chose only one chain per deposit, so the terms chain, deposit and structure are used in this publication interchangeably.

### 2.3 Identification of P450 domains

Due to low sequence similarity between P450 superfamily members, we frequently observe that simple sequence-based searches lack sufficient sensitivity, which carries a substantial risk of missing relevant proteins. To identify all potential P450 proteins in the PDB, we applied a two-step procedure. First, we retrieved all previously annotated enzymes belonging to this superfamily, as well as proteins that can be readily identified based on established objective criteria. Second, we scanned the PDB using structural representative P450 structures to detect additional candidates.

#### 2.3.1 Initial search based on keywords and P450atlas classification

In the first step, we performed a search for deposits containing phrases CYP or P450 and containing heme within the structure. Some P450s or P450-like proteins contain a different cofactor (e.g. a similar molecule, but with the iron atom replaced by a different element) or lack these keywords in the description, but such instances are rare and were easily found in the second part of the search. Sequences from all identified deposits were submitted to the P450atlas subfamily assignment server to determine whether they truly correspond to P450 enzymes and to obtain their initial CYPid annotations. As a results, we have identified 1358 cytochrome-containing deposits, which were manually verified and used in further search.

#### 2.3.2 Search using structural similarity

In order to find any P450 deposits missed during the previous step, we also conducted a structure-based search, computing structural alignments between every chain in the PDB and a set of representative P450 templates. We extracted a total of unique 466,228 polymer chains from the PDB database. To limit the computational effort, we excluded chains that, for objective reasons, cannot be P450s — namely, those shorter than 200 amino acids or containing fewer than five *α*-helices. Applying these filters sequentially reduced the dataset to 219,802 chains (helix criterion) and 223,363 chains (length criterion). After applying both criteria simultaneously, we were left with 182,844 chains. All previously characterized P450 enzymes contain at least 11 *α*-helices and 328 residues, which indicates that our choice of cut-offs is extremely unlikely to exclude any genuine P450s.

To further reduce the computational effort, we have decided to select only several P450 representatives from the previously found chains. To obtain them, we have performed the complete-link clustering, with distance defined as *d* = (1 − TM score), where TM score is defined as in [24] and the clustering cut-off distance set to *d* = 0.3. This way, we have gotten three clusters and, for each of them, the representative was chosen as the structure, for which the largest distance to any other structure in the same cluster was the lowest. The three representatives were the A chains of 3EL3, 7WEX: and the 7TLO.

We then performed the structural alignment of each of the 182,844 pre-selected chains with each of the three P450 representatives. This let us identify another 92 deposits with P450s, which had not been found at the previous stage because of a lacking deposit description or an alternative heme variant. The total number of results after the second stage was 1450.

### 2.4 Family and subfamily assignment

For each deposit, we searched through its description and, if available, the abstract of the related publication, to find the corresponding CYPid. Simultaneously we ran the P450atlas family assignment server for all of them, obtaining the best matches for all of them. Finally, we manually checked the sequences against the database and manually assigned the proper CYPids.

### 2.5 Table update procedure

We have automated the described structure search procedure to create a straightforward pipeline enabling regular updates to the publicly available P450 deposit table with little need for manual curation. It is going to be run every quarter to keep the structural data hosted on the p450atlas website up to date. The pipeline consists of (i) identifying all new protein entities deposited into PDB since the previous update, (ii) filtering out the proteins with less than 100 residues, (iv) selecting the most complete chain for each entity, (v) running TMalign between those chains and the three P450 representatives and rejecting those chains for which the highest TM-score was below 0.6 and finally (vi) running the p450atlas subfamily assignment and (vii) manually verifying any unobvious cases.

Since the original search was performed on a database from July 28th 2024, we have tested this procedure to include deposits from between that day and beginning of the year 2026. We have found 63 structures deposited to the PDB within the 18 months of that period, making 1513 total P450-containing deposits, as of January 1st 2026.

## 3 Results and Discussion

In total, we found 1513 deposits in PDB, each containing a distinct P450 enzyme structure. However, since there are multiple deposits for some proteins, there were only 674 unique sequences. The first stage of our search identified 1358 deposits, the structure-based search found 92 more and the update procedure 63. Some of the results found with the structural search were missed in the first stage because they had another heme variant than the expected Heme B (Protoporphyrin IX with iron atom). Others lacked the pre selected keywords (for example the deposit authors only provided the common name).

### 3.1 Similarity between P450 proteins

While P450 enzymes exhibit very high sequence diversity, with pairwise sequence identity dropping below 20%, they all retain distinct structural features, and their structural similarity — quantified using metrics such as the TM-score — remains consistently high. We illustrate this in Figure 2, which shows the distributions of sequence identity and TM-score for all possible pairs of P450 structures obtained thus far. Notably, while typical sequence identities range anywhere from 14% to 60%, with an additional peak near 100% corresponding to multiple deposits of the same CYP, TM-scores are predominantly above 0.7, even for sequence identities below 20%. The TM-score parameter is used to quantify the structural similarity between two protein structures. It ranges from 0 to 1, where values closer to 1 indicate highly similar folds, while scores below 0.17 typically correspond to unrelated structures. Result above 0.5 indicates that the two proteins share the same overall fold. Our results therefore demonstrates that even the most divergent cytochromes P450 share substantial structural similarity, and when considering the maximum TM-score between a query and multiple representatives — rather than the minimum — the effective cutoff can be set even higher. As visible in Figure 2, the mode of the histogram, and thus the most probable case, corresponds to a pair of proteins with 22% sequence identity and a TM-score of 0.82. An example of such a pair is given in Figure 3. Interestingly, for each structure in our dataset, there is at least one other chain with which the sequence identity is below 25% while the TM-score is above 0.8, which further demonstrates how strongly the structure is preserved in the P450 superfamily.

**Figure 2:**
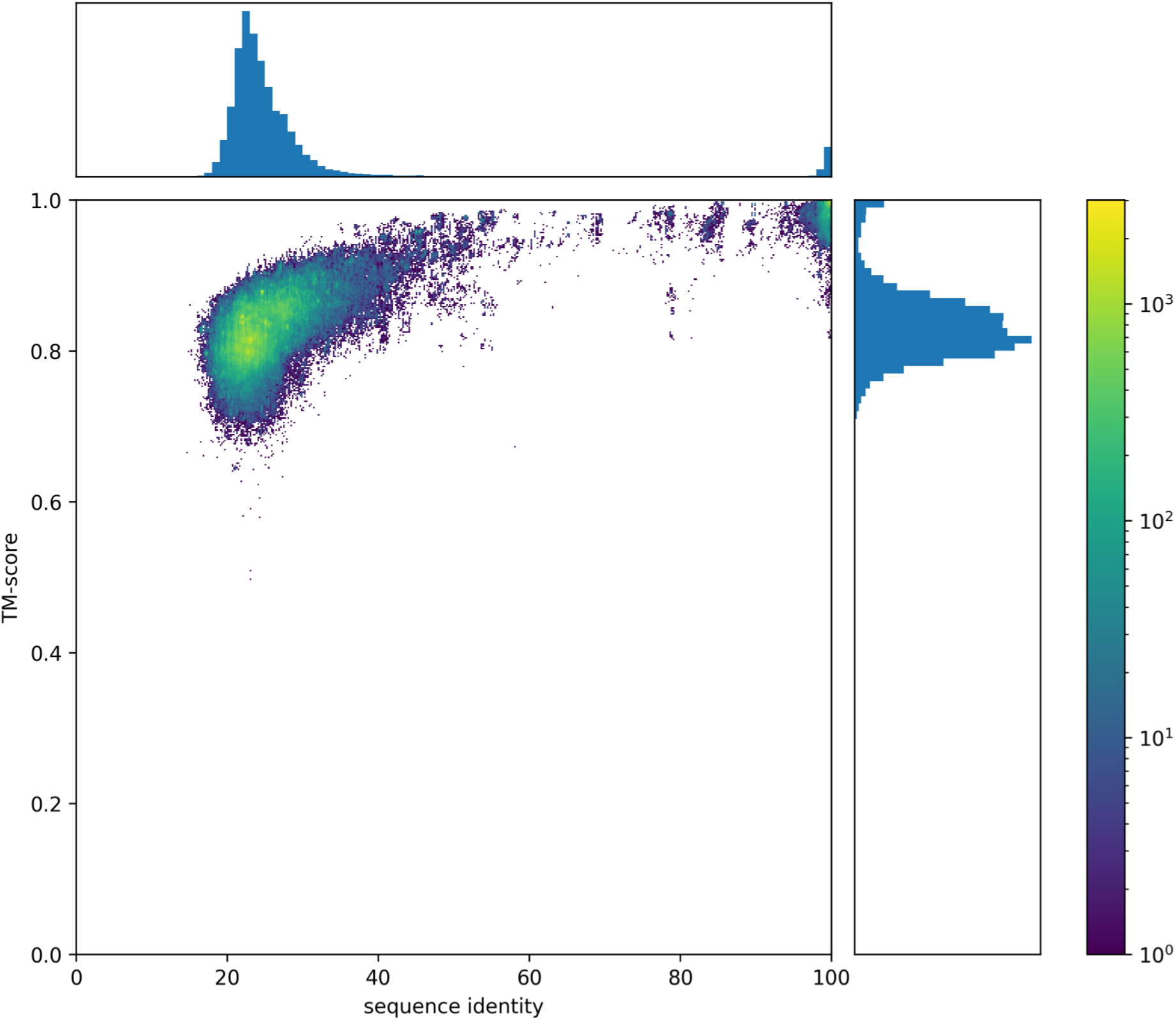
Distribution of the values of TM-score and sequence identity for all possible protein pairs in the structures dataset after the initial keyword and assignment server search. The white spaces on the 2-dimensional histogram mean no observations. TM-scores are typically very high (above 0.7) even for the low sequence identity

**Figure 3:**
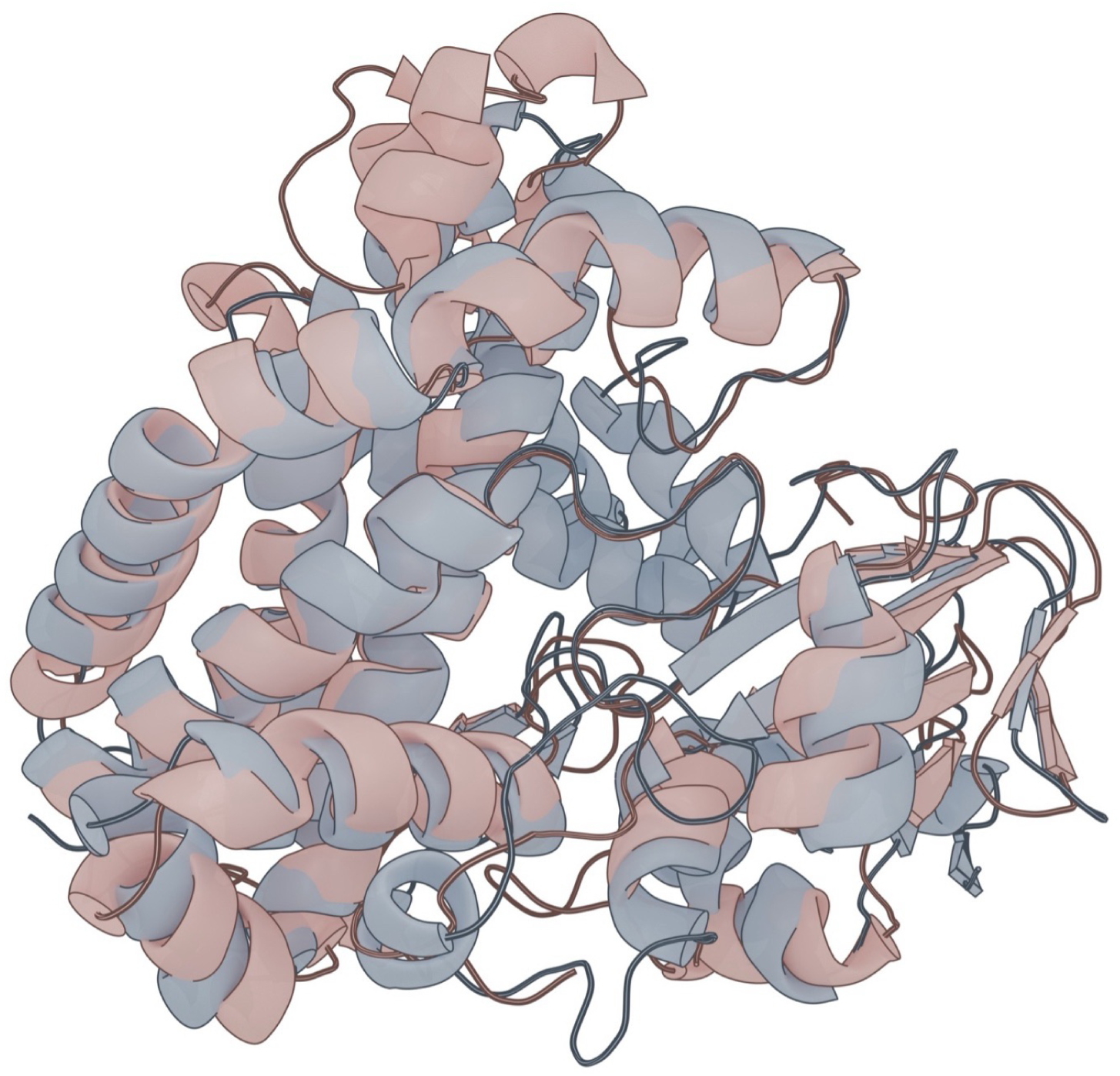
A typical example of two P450 proteins that exhibit high structural similarity despite low sequence identity. CYP175A (chain A of 1N97, brown) and CYP165E (chain A of 7TTP, blue) are identical in 22.9% by sequence while TM-score value is 0.82. The crmsd value over 369 aligned residues is 3.37 Å.

### 3.2 Classification by the deposit authors

We carefully inspected the deposit information and the corresponding publication abstracts for all the structures found, to identify the CYPid provided by the deposit authors. If no CYPid was given, but the family or subfamily was otherwise indicated (e.g. *P450-102* meaning family CYP102) we treated it the same as if the CYPid had been given directly. We treated all the common name notations (like *P450-CAM* and *P450(cam)*) as equivalent and case-insensitive and gathered them for the found deposits.

The CYPids and common names could be found in several different places in the deposit, primarily the deposit title, gene name and entity name. Sometimes none of these fields contained the CYPid, but the corresponding article abstract did. Many deposits were poorly labeled. Frequently the deposit label, gene name and the polymer entity name were inconsistent, making manual inspection of the deposits necessary. The entity name was often incorrect — listing some remote homolog — or ambiguous — consisting of multiple CYPids or names. We found that misclassification was rare, but lack of specificity and usage of non-standardized nomenclature were very common. The examples of the prior are providing only the family without the subfamily, as well as calling the protein *CYPxxx-homolog*. The latter typically means usage of some *common name* instead of CYPid or some obsolete or non-standard notation. In some cases, beside the standard CYPid or the common name, an obsolete CYPid equivalent was present, where a roman numeral was used to denote the family, e.g. *P450-LIA* instead of CYP51A. Such examples are not counted in our statistics or listed in the table.

If there was any ambiguity in the classification in the deposits (e.g. multiple CYPids or aliases were provided), we assumed the interpretation more favorable to the deposit authors. In total, out of 1513 deposits, 905 had the correct family and subfamily provided, 287 had the correct family but no subfamily, 284 had no family, but some alias (common name) was given, 16 had no family information and in one case there was no indication at all, that the structure was of a P450 enzyme. In 19 cases the family given by authors was correct, but the subfamily was wrong and in 2 cases the family was incorrect. In one case there was no information whatsoever, that the structure was of Cytochrome P450. These results are summarized in Figure 4.

**Figure 4:**
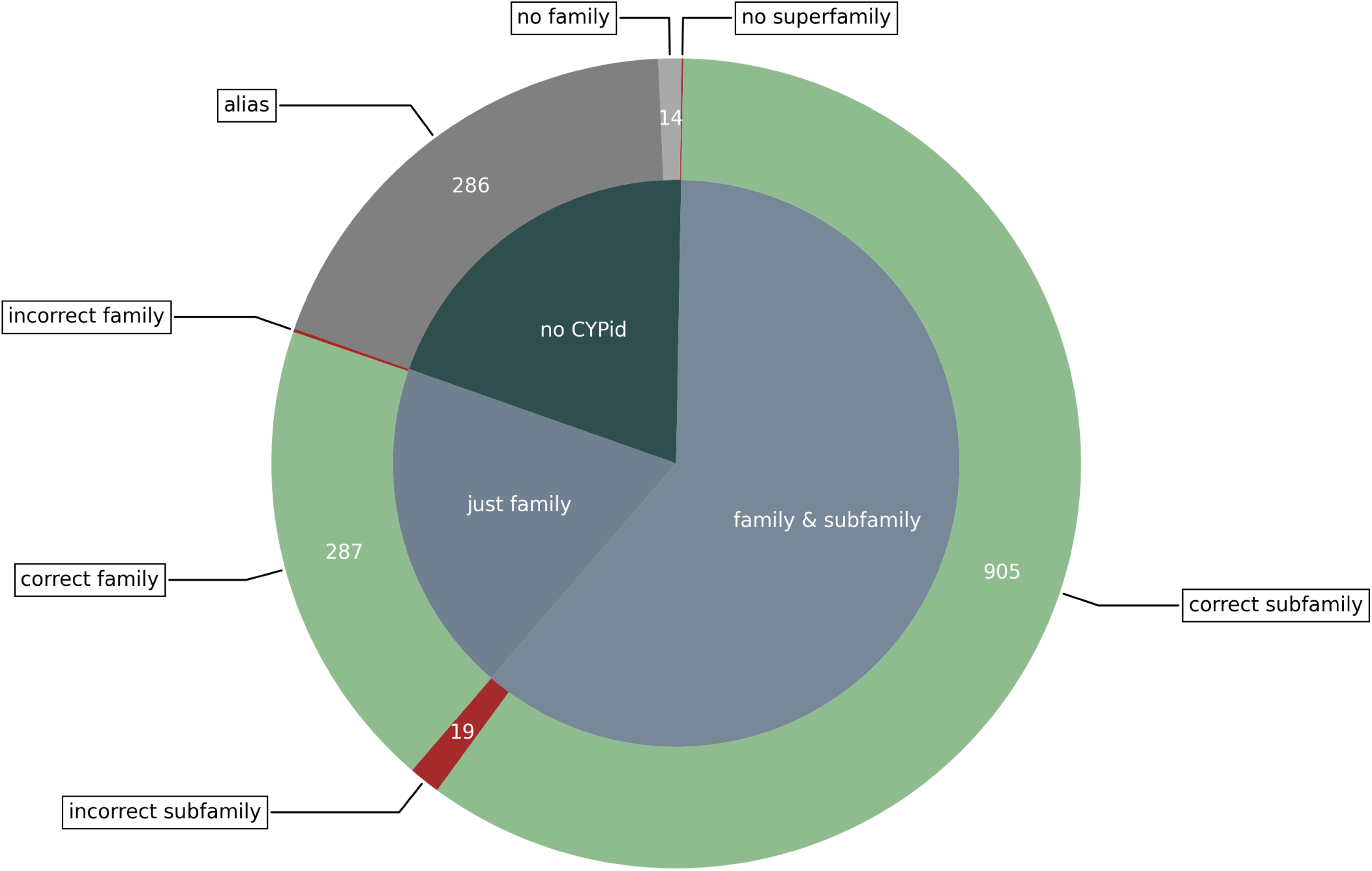
Precision and correctness of families and subfamilies as given in the publication or the structure deposit information; *incorrect subfamily* refers to the cases where the family was properly assigned, but the subfamily was incorrect, *no subfamily* refers to cases where family was properly assigned, but subfamily wasn’t provided at all.

Two deposits for which the wrong family had been assigned are 5XW2 and 4E2P, both of which are categorized as *P450 107B1* in the deposit file, while the proper CYPid for them is CYP113D2. They are primarily referred to as *HmtN* or *Himastatin hydroxylase* by the deposit authors. One case in which not only no family, subfamily or a common name was given, but there was also no mention of terms like *P450* or *Cytochrome* was 8D8P, a CYP152T9 enzyme from *Rothia nasimurium*, in complex with the palmitic acid. In the deposit information, the protein is referred to as a *fatty acid decarboxylase* or as an *oxidoreductase*.

### 3.3 Common names in the deposits

Common names were found in 980 deposits, including some, for which no family was indicated. Figure 5 shows the ones with at least 20 corresponding deposits.

**Figure 5:**
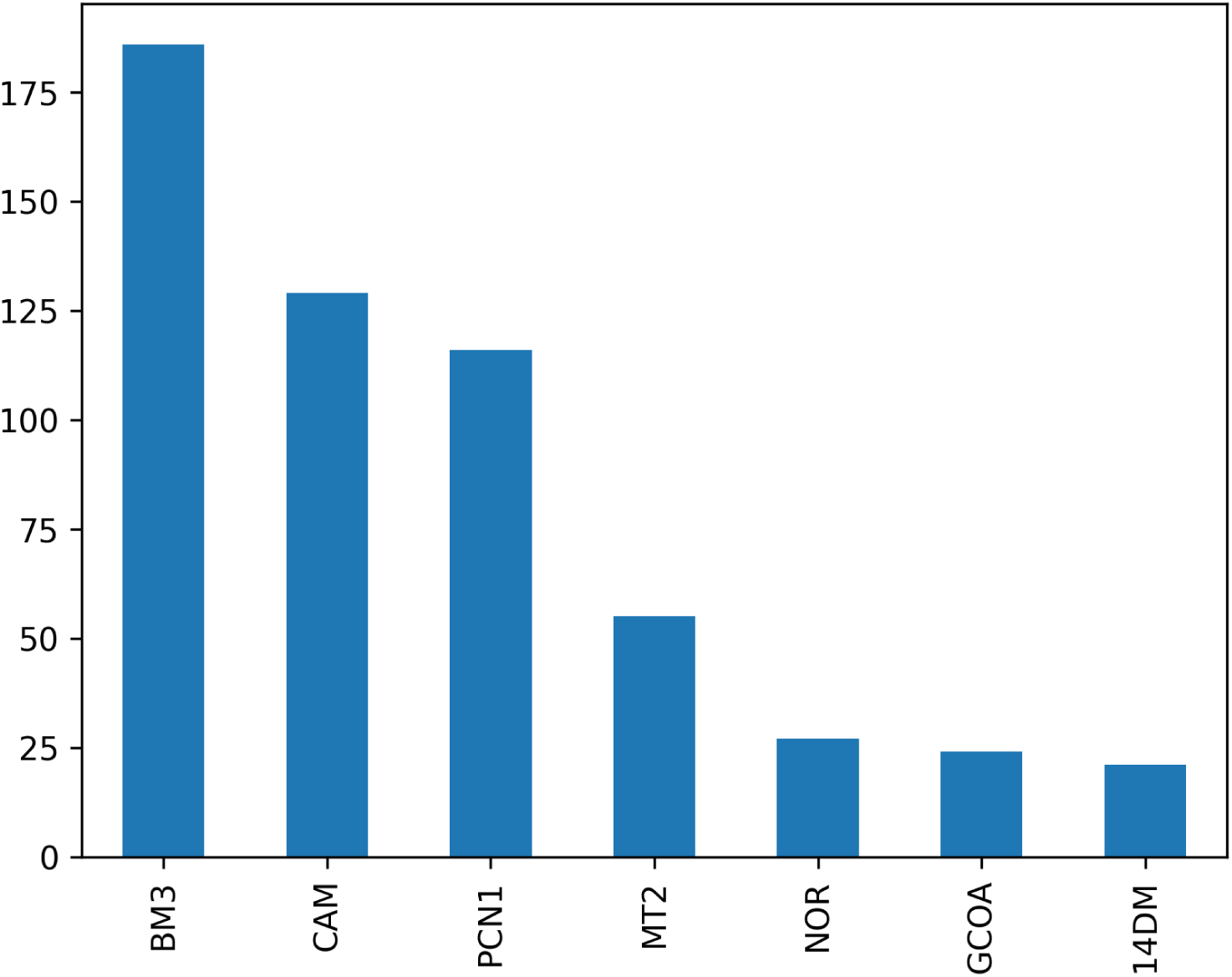
The most frequent common names in our dataset. Since case and punctuation are not always consistent for some of them, they are all provided with no punctuation and in uppercase

The first place is occupied by *P450 BM3* (originally *P450_BM-3_*), with 186 corresponding structures, which refers to CYP102A1, a fatty acid hydroxylase. It comes from the fact that it was the third cytochrome isolated from *Priesta megaterium* [25], which had originally been called *Bacillus megaterium* (BM) but was renamed around 2020 [26]. It was the second cytochrome, for which a structure had been obtained (deposit 2HPD) [27].

The second most popular common name *P450 CAM* (originally *P 450CAM*) is present in 129 deposits and refers to the CYP101A1, which is responsible for hydroxylation of camphor. The name comes from the substrate. It was the first, and until the publication of P450 BM3 structure the only, cytochrome for which an experimental structure was available (deposit 2CPP) [28]. CYP101 is the second most popular family in our dataset, with 144 corresponding deposits.

The third place is occupied by *P450-PCN1* (originally *P450PCN1*), meaning the first pregnenolone 16 alpha-carbonitrile (PCN)-induced P-450. In the presented dataset it refers exclusively to human CYP3A4, although it originally referred to rat CYP3A1 [29, 30]. In literature CYP3A4 is sometimes called *human P450-PCN1*, differentiating it from the rat enzyme [31]. It is also known as *HLp* (human liver protein) and *NF-25* (Nifedipine oxidase clone 25) [32] and both these names are present in CYP3A4 deposits, next to PCN1.

The names *BM3*, *CAM* and *PCN1* all have over 100 corresponding deposits. The next most common name with 55 corresponding deposits is *P450-MT2* (originally P*450 MT2*). It denotes CYP121A1, a known drug target and comes from the host organism Mycobacterium Tuberculosis [33]. No other common name has over thirty corresponding deposits.

Apart from the already mentioned CYP3A4, several other P450s have multiple aliases. For instance, its ortholog, CYP3A7 is usually referred to either as *P450-HLp2* (Human Liver p450 nr 2) or *HFLA* (Human Fetal Liver enzyme a) [34]. CYP55A1, typically called *P450nor*, is also sometimes called *P450 DNIR*, and CYP11B2 is known both as *P450aldo* (aldosterone synthase) and *P-450C18* (steroid 18-hydroxylase)[35]. For each entry with multiple aliases, we only provide one in the results table (e.g. *P450-PCN1* for CYP3A4 and *P450nor* for CYP55A1).

The inverse issue also exists, i.e. a single alias is sometimes used for multiple enzymes. As an example, *P450scc* (side-chain cleavage enzyme) typically refers to CYP11A1, a key human P450 responsible for conversion of cholesterol to pregnenolone [36], but the authors of the 9WAT deposit use it also in reference to CYP204A1, which is a bacterial side-chain cleaveage enzyme [37].

All common names are listed in the deposit table in the supplementary materials.

### 3.4 Classification by the assignment server and new subfamilies

Since P450atlas runs the family assignment for sequences rather than the structures and the assignment is fully deterministic, we discuss the statistics for unique sequences, rather than the deposits.

For a given query sequence, the P450atlas server first uses Hidden Markov Model profiles to pre-select the best-fitting Cytochrome subfamilies, followed by the global alignment with the representative amino acid sequences of said subfamilies. If the identity between the query and the most similar representative sequence is above the 55% threshold, the server recommends the subfamily. If it is between 40% and 55%, only the family is suggested and if it is below 40% no family is assigned. Nevertheless, the pre-selected families along with their representatives and the corresponding sequence identities are always listed. Importantly, especially for the larger subfamilies, not all members are selected for alignment, meaning that for some queries the sequence identity provided by P450atlas is underestimated.

In case of the discussed dataset, for 14 different sequences (32 deposits in total), the best sequence identity provided by the server was below the 55% threshold, and no subfamily was explicitly recommended. However, after a careful evaluation by a human expert, considering all sequences (not just the representatives) and the evolutionary context, it was determined that for all but five of those sequences (7 deposits), the best hit from P450Atlas was the correct subfamily. The remaining sequences were placed in new subfamilies: the sequence from deposits 8U19, 8U1I and 8U09 in CYP255D, sequences from 9DUJ and 9EBY in CYP1251G and sequences from 9KPP and 9KPU in CYP107PW.

After the expert evaluation, 4 additional sequences, for which the sequence identity with the closest representative was above 55%, were assigned to two new subfamilies (CYP165F and CYP152AX). In total, 5 new subfamilies were created to which 9 different sequences from 14 deposits were assigned. All deposits with new families are listed in Table 1 along with the best fitting previously existing family according to P450atlas.

**Table 1:** Deposits (grouped by sequence) with new subfamilies and the best matches according to the assignment server.

| Deposits | Best match according to P450atlas | New subfamily |
| --- | --- | --- |
| 7TTO, 7TTP, 7TTQ | CYP165E | CYP165F |
| 6M7L | CYP165E | CYP165F |
| 7TTA | CYP165E | CYP165F |
| 8W1K, 8W1J | CYP152T | CYP152AX |
| 8U19, 8U1I, 8U09 | CYP255A | CYP255D |
| 9KPP | CYP107G | CYP107PW |
| 9KPU | CYP107G | CYP107PW |
| 9DUJ | CYP1251A | CYP1251G |
| 9EBY | CYP1251A | CYP1251G |

### 3.5 Represented families

The 1513 P450 proteins found in the Protein Data Bank come from 86 different families. However, the dataset is not particularly diverse, with 62.39% deposits belonging to just 8 largest families. Five of them have over 100 deposits and three more have over 50. Moreover, 51 families are represented by just a single CYPid. While all the largest families have more than one CYPid in the dataset, most deposits usually still belong to a single ortholog group. For instance, CYP102 is the largest family in the dataset with 197 corresponding deposits, 193 of which are CYP102A1, also known as P450-BM3. CYP101, the second largest family, has 146 deposits, 129 of which are CYP101A1 (P450-CAM). Similar results are observed for CYP3 with CYP3A4, for CYP199 with CYP199A4 and CYP121 with CYP121A1. More diverse are CYP51, CYP2 and CYP107. The pie chart with the largest families and ortholog groups is presented in Figure 6.

**Figure 6:**
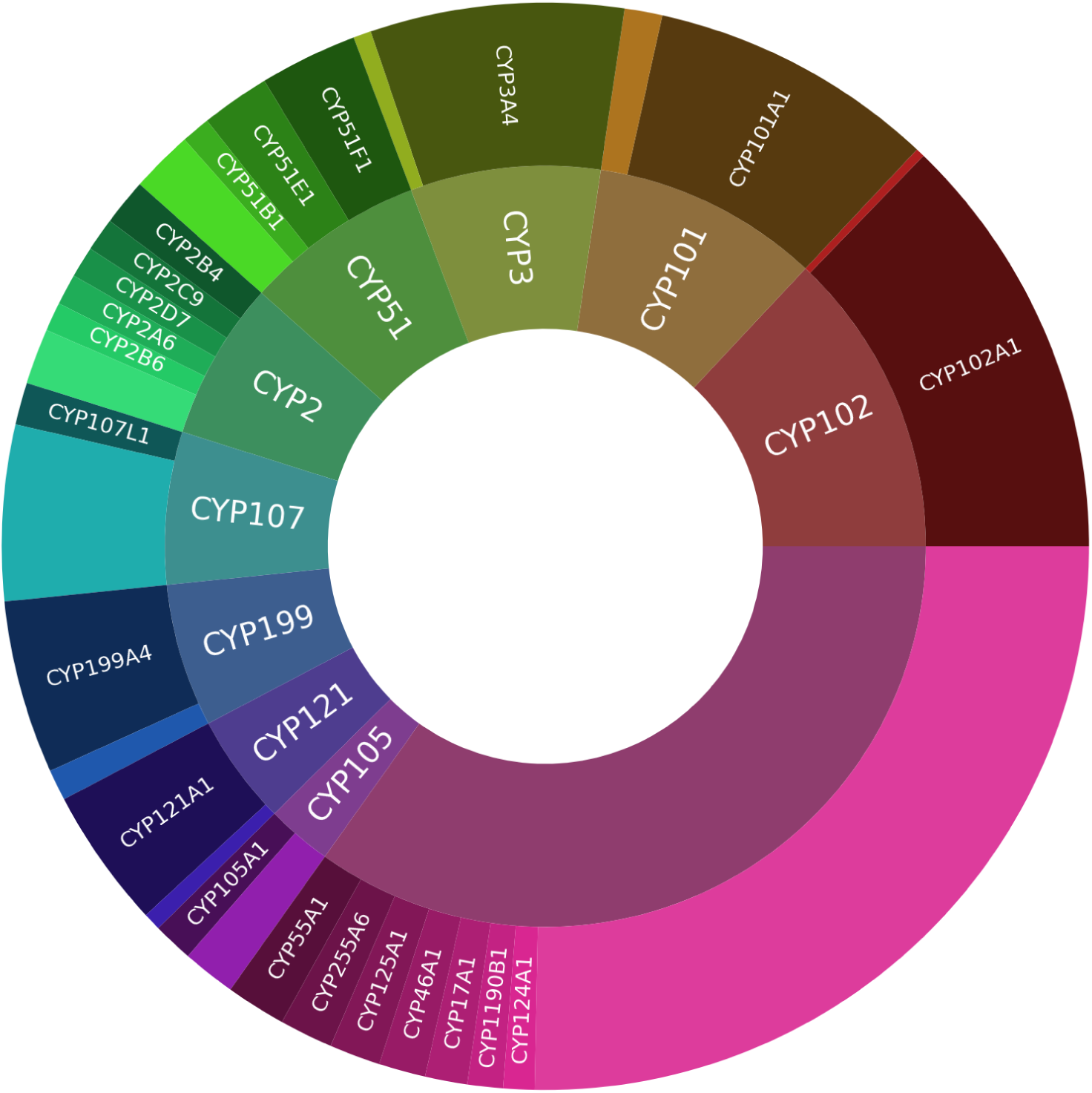
P450s represented in the dataset, grouped by family. For readability only the families with at least 30 and ortholog groups with at least 12 corresponding deposits are mentioned explicitly. The unlabeled pieces in the outer circles represent ortholog groups with less than 12 deposits and in the inner circle they represent the families with less than 30 deposits

The large number of experimental structures for these best-represented families reflects their importance. CYP102 are naturally self-sufficient P450s, since the heme domain and reductase domain are fused into a single chain, eliminating the need for separate redox partners. As mentioned in the section regarding *common names*, CYP102A1 (P450-BM3) was the second P450 to be crystallized and has since served as a model P450, highly explored for biotechnological applications, thanks to its self-sufficiency and high catalytic turnover [38]. Before the release of the first CYP102 in 1993, CYP101A1 was the only available P450 structure. Thus, it was extensively used as a model system for P450 research: understanding the P450 structure-functional relationships, heme-binding, and catalytic mechanisms [39] It has also been engineered for various biotechnological applications. CYP2 and CYP3 families’ popularity stems largely from their role in drug metabolism [40]. The most common of them, CYP3A4, also mentioned in the *common names* section and known as *P450-PCN1*, *HLp* and *NF-25*, is responsible for metabolism of over half of all prescribed drugs and thus remains a crucial object of research [41]. CYP51 is a well-known target for azole drugs important in treatment of fungal infections and has therefore been subject of structure-function analysis to not only assess azole drug interactions but also refine or identify its new inhibitors [42, 43, 44]. CYP107 family members, on the other hand, are biotechnologically valuable, as they are involved in the generation of beneficial secondary metabolites [45].

### 3.6 Alternative heme variants

One of the most basic attributes of Cytochrome P450s is the cysteine-bound heme b, which in PDB is assigned the HEM code. However, as mentioned earlier, multiple CYP deposits have no HEM molecule, but rather some other cofactor. We list all those alternative cofactors in 2.

By far, the most common one was HEC. In PDB it refers to heme C, characterized by thioether bonds to cysteines in place of the typical vinyl groups. In case of the discussed deposits, however, the code seems to be assigned erroneously. The article corresponding to 7ABB deposit, for instance, explicitly refers to the heme as heme b, despite it being called heme C in the deposit [46] and none of the deposits with HEC possess said thioether linkages.

The second most popular MI9 is Oxomolybdenum Mesoporphyrin IX, a heme analog with oxygen bound molybdenum atom in place of Iron. It mimics the highly reactive oxoferryl heme species (Compound 1) but is itself stable and unreactive. As such, it is used, e.g. in 7WY1 deposit, to capture a particular shortlived stage of the P450 catalytic cycle [47, 48].

**Table 2:** Alternative heme cofactors and the corresponding PDB deposit codes.

| HEME | PDB-IDs |
| --- | --- |
| HEC | 7ABB,1OG5,1OG2,7NQM,7NQO,7ANT,2H7S,6YCT,7ABA,6XZ8,6XZ9 |
| MI9 | 7WY1,7W9D,7W9J,6JMH,7WY3,7WY2 |
| HEB | 4L36,8C9I,8ABS,8ABR |
| MNH | 5ZIS,5ZLH,6JZS,7WG0 |
| MNR | 2FEU,2FER,2FE6 |
| FDE | 5JQV,5JQU |
| PP9 | 6G71,5E9Z |
| 7HE | 2Z97 |
| BW9 | 6JMW |
| COH | 6JVC |
| CV0 | 6K24 |
| RUR | 6K9S |
| HIR | 7UOR |
| MH0 | 8E5J |
| 6HE | 2ZAW |

PP9 is protoporphyrin IX without the iron atom. However, both deposits listed with that cofactor do actually have the iron atom, but listed separately from the cofactor. Ten deposits listed here come from an experiment where CYP102A1 was crystallized with alternative hemes, in which the Iron atom had been replaced with some other element; in particular, in BW9 iron is replaced by chromium, in COH by Cobalt, in CV0 by Rhodium and in RUR by Rhutenium [49]. In both MNH and MNR Iron is replaced by Manganese and finally HIR is methyliridium(III) mesoporphyrin.

## 4 Conclusion

Cytochrome P450 monooxygenases (CYPs/P450s) have been in focus for the past six decades due to their extensive applications across various fields of biology, primarily because of their catalytic diversity, which enables multiple enzymatic reactions with chemo-, regio-, and stereo-selectivity. Due to their potential biotechnological applications, many P450s have been crystallized, and researchers have deposited these structures under various names, often without adhering to the P450 classification and nomenclature guidelines. The frequently used *common names* often lack specificity and are inconsistently typed. Both these names and CYPids can be found in multiple different fields in the PDB deposits, making any automated search very difficult.

In this study, using a combination of bioinformatic tools, we have performed an extensive search and analysis of P450 structures in the PDB. Finally, we have assigned a correct CYPid (family and subfamily) for all the identified enzymes, enabling researchers to identify and cite these P450s properly. The P450Atlas server was found to be accurate in assigning P450 families and subfamilies.

A table listing all P450 deposits with their family and subfamily information is provided in this article (as a supplementary file) and is also available online on the P450atlas.org website, serving as a valuable resource for P450 researchers. Additionally, all P450 sequences deposited in PDB have been incorporated into the P450atlas database to enhance the accuracy of the CYPid assignment procedure implemented on this server.

## Supporting information

XLSX table with all P450 deposits identified by this study

## References

[1] D. Gront, K. Syed, and D. R. Nelson, “Exploring P450 superfamily diversity with P450ATLAS - Online tool for automated subfamily assignment,” Protein Science, vol. 34, no. 3, p. e70057, Mar. 2025. [Online]. Available: https://onlinelibrary.wiley.com/doi/10.1002/pro.70057

[2] D. Garfinkel, “Studies on pig liver microsomes. I. Enzymic and pigment composition of different microsomal fractions,” Archives of Biochemistry and Biophysics, vol. 77, no. 2, pp. 493–509, Oct. 1958. [Online]. Available: https://www.sciencedirect.com/science/article/pii/000398615890095X

[3] M. Klingenberg, “Pigments of rat liver microsomes,” Archives of Biochemistry and Biophysics, vol. 75, no. 2, pp. 376–386, Jun. 1958. [Online]. Available: https://www.sciencedirect.com/science/article/pii/0003986158904363

[4] T. Omura and R. Sato, “A New Cytochrome in Liver Microsomes,” Journal of Biological Chemistry, vol. 237, no. 4, pp. PC1375–PC1376, Apr. 1962. [Online]. Available: https://www.sciencedirect.com/science/article/pii/S0021925818603382

[5] R. E. White and M. J. Coon, “Oxygen Activation by Cytochrome P-4501,” Annual Review of Biochemistry, vol. 49, no. Volume 49, 1980, pp. 315–356, Jul. 1980. [Online]. Available: https://www.annualreviews.org/content/journals/10.1146/annurev.bi.49.070180.001531

[6] F. P. Guengerich, “Cytochrome P450 research and *The Journal of Biological Chemistry*,” Journal of Biological Chemistry, vol. 294, no. 5, pp. 1671–1680, Feb. 2019. [Online]. Available: https://www.sciencedirect.com/science/article/pii/S0021925820364802

[7] M. Katagiri, B. N. Ganguli, and I. C. Gunsalus, “A soluble cytochrome P-450 functional in methylene hydroxylation.” The Journal of biological chemistry, vol. 243, no. 12, pp. 3543–3546, 1968.

[8] I. C. Gunsalus and S. G. Sligar, “Oxygen Reduction by the P450 Monoxygenase Systems,” in Advances in Enzymology and Related Areas of Molecular Biology. John Wiley & Sons, Ltd, 1979, pp. 1–44. [Online]. Available: https://onlinelibrary.wiley.com/doi/abs/10.1002/9780470122921.ch1

[9] T. L. Poulos, B. C. Finzel, and A. J. Howard, “High-resolution crystal structure of cytochrome P450cam,” Journal of Molecular Biology, vol. 195, no. 3, pp. 687–700, Jun. 1987. [Online]. Available: https://www.sciencedirect.com/science/article/pii/0022283687901902

[10] D. Gront, K. Syed, and D. R. Nelson, “Exploring P450 superfamily diversity with P450ATLAS - Online tool for automated subfamily assignment,” Protein Science, vol. 34, no. 3, p. e70057, Mar. 2025. [Online]. Available: https://onlinelibrary.wiley.com/doi/10.1002/pro.70057

[11] D. W. Nebert, M. Adesnik, M. J. Coon, R. W. Estabrook, F. J. Gonzales, F. P. Guengerich, I. C. Gunsalus, E. F. Johnson, B. Kemper, W. Levin, I. R. Phillips, R. Sato, and M. Waterman, “The P450 Gene Superfamily: Recommended Nomenclature,” DNA, vol. 6, no. 1, pp. 1–11, Feb. 1987.

[12] D. W. Nebert, D. R. Nelson, M. Adesnik, M. J. Coon, R. Estabrook, F. J. Gonzales, F. P. Guengerich, I. C. Gunslaus, E. F. Johnson, B. Kemper, W. Levin, I. R. Phillips, R. Sato, and M. Waterman, “The P450 Superfamily: Updated Listing of All Genes and Recommended Nomenclature for the Chromosomal Loci,” DNA, vol. 8, no. 1, pp. 1–13, Jan. 1989.

[13] D. W. Nebert, D. R. Nelson, M. J. Coon, R. W. Estabrook, R. Feyereisen, YOSHIAKI. Fujii-Kuriyama, F. J. Gonzales, F. P. Guengerich, I. C. Gunslaus, E. F. Johnson, J. C. Loper, R. Sato, M. Waterman, and D. J. Waxman, “The P450 Superfamily: Update on New Sequences, Gene Mapping, and Recommended Nomenclature,” DNA and Cell Biology, vol. 10, no. 1, pp. 1–14, Jan. 1991.

[14] D. R. Nelson, T. Kamataki, D. J. Waxman, F. P. Guengerich, R. W. Estabrook, R. Feyereisen, F. J. Gonzales, M. J. Coon, I. C. Gunslaus, O. Gotoh, K. Okuda, and D. W. Nebert, “The P450 Superfamily: Update on New Sequences, Gene Mapping, Accession Numbers, Early Trivial Names of Enzymes, and Nomenclature,” DNA and Cell Biology, vol. 12, no. 1, pp. 1–51, Jan. 1993.

[15] D. R. Nelson, “The Cytochrome P450 Homepage,” Human Genomics, vol. 4, no. 1, p. 59, Dec. 2009.

[16] D. R. Nelson, “Cytochrome P450 Nomenclature, 2004,” in Cytochrome P450 Protocols. New Jersey: Humana Press, 2004, pp. 1–10.

[17] H. M. Berman, J. Westbrook, Z. Feng, G. Gilliland, T. N. Bhat, H. Weissig, I. N. Shindyalov, and P. E. Bourne, “The Protein Data Bank,” Nucleic Acids Research, vol. 28, no. 1, pp. 235–242, Jan. 2000. [Online]. Available: 10.1093/nar/28.1.235

[18] H. M. Berman and S. K. Burley, “Protein Data Bank (PDB): Fifty-three years young and having a transformative impact on science and society,” Quarterly Reviews of Biophysics, vol. 58, p. e9, Jan. 2025. [Online]. Available: https://www.cambridge.org/core/journals/quarterly-reviews-of-biophysics/article/protein-data-bank-pdb-fiftythree-years-young-and-having-a-transformative-impact-on-science-and-society/36E04BF6CB3DE157DF3E09EFD8EC47DC

[19] J. M. Macnar, N. A. Szulc, J. D. Kryś, A. E. Badaczewska-Dawid, and D. Gront, “BioShell 3.0: Library for Processing Structural Biology Data,” Biomolecules, vol. 10, no. 3, p. 461, Mar. 2020.

[20] “PyMOL Open Source.” [Online]. Available: https://github.com/schrodinger/pymol-open-source

[21] Y. Zhang, “TM-align: A protein structure alignment algorithm based on the TM-score,” Nucleic Acids Research, vol. 33, no. 7, pp. 2302–2309, Apr. 2005.

[22] “Blender - a 3D modelling and rendering package,” Blender Foundation. [Online]. Available: https://www.blender.org/

[23] J. D. Kryś and D. Gront, “VisuaLife: Library for interactive visualization in rich web applications,” Bioinformatics, vol. 37, no. 20, pp. 3662–3663, Oct. 2021.

[24] Y. Zhang and J. Skolnick, “Scoring function for automated assessment of protein structure template quality,” Proteins: Structure, Function, and Bioinformatics, vol. 57, no. 4, pp. 702–710, Dec. 2004. [Online]. Available: https://onlinelibrary.wiley.com/doi/10.1002/prot.20264

[25] C. J. C. Whitehouse, S. G. Bell, and L.-L. Wong, “P450BM3 (CYP102A1): Connecting the dots,” Chemical Society Reviews, vol. 41, no. 3, pp. 1218–1260, Jan. 2012. [Online]. Available: https://pubs.rsc.org/en/content/articlelanding/2012/cs/c1cs15192d

[26] R. S. Gupta, S. Patel, N. Saini, and S. Chen, “Robust demarcation of 17 distinct Bacillus species clades, proposed as novel Bacillaceae genera, by phylogenomics and comparative genomic analyses: Description of Robertmurraya kyonggiensis sp. nov. and proposal for an emended genus Bacillus limiting it only to the members of the Subtilis and Cereus clades of species,” International Journal of Systematic and Evolutionary Microbiology, vol. 70, no. 11, pp. 5753–5798, 2020. [Online]. Available: https://www.microbiologyresearch.org/content/journal/ijsem/10.1099/ijsem.0.004475

[27] K. G. Ravichandran, S. S. Boddupalli, C. A. Hasermann, J. A. Peterson, and J. Deisenhofer, “Crystal Structure of Hemoprotein Domain of P450BM-3, a Prototype for Microsomal P450’s,” Science, vol. 261, no. 5122, pp. 731–736, Aug. 1993.

[28] T. L. Poulos, B. C. Finzel, and A. J. Howard, “High-resolution crystal structure of cytochrome P450cam,” Journal of Molecular Biology, vol. 195, no. 3, pp. 687–700, Jun. 1987.

[29] F. J. Gonzalez, D. W. Nebert, J. P. Hardwick, and C. B. Kasper, “Complete cDNA and protein sequence of a pregnenolone 16 alpha-carbonitrile-induced cytochrome P-450. A representative of a new gene family.” Journal of Biological Chemistry, vol. 260, no. 12, pp. 7435–7441, Jun. 1985. [Online]. Available: https://linkinghub.elsevier.com/retrieve/pii/S0021925817396266

[30] F. J. Gonzalez, B. J. Song, and J. P. Hardwick, “Pregnenolone 16 alpha-carbonitrile-inducible P-450 gene family: Gene conversion and differential regulation.” Molecular and Cellular Biology, vol. 6, no. 8, pp. 2969–2976, Aug. 1986. [Online]. Available: https://pmc.ncbi.nlm.nih.gov/articles/PMC367867/

[31] F. J. Gonzalez, B. J. Schmid, M. Umeno, O. W. Mcbride, J. P. Hardwick, U. A. Meyer, H. V. Gelboin, and J. R. Idle, “Human P450PCN1: Sequence, Chromosome Localization, and Direct Evidence through cDNA Expression That P450PCN1 Is Nifedipine Oxidase,” DNA, vol. 7, no. 2, pp. 79–86, Mar. 1988. [Online]. Available: https://journals.sagepub.com/doi/full/10.1089/dna.1988.7.79

[32] “UniProtKB P08684 · CP3A4 HUMAN.” [Online]. Available: https://www.uniprot.org/uniprotkb/P08684/entry

[33] T. Padayachee, D. C. Lamb, D. R. Nelson, and K. Syed, “Structure-Function Analysis of the Essential Mycobacterium tuberculosis P450 Drug Target, CYP121A1,” International Journal of Molecular Sciences, vol. 25, no. 9, p. 4886, Apr. 2024.

[34] “UniProtKB P24462 · CP3A7 HUMAN.” [Online]. Available: https://www.uniprot.org/uniprotkb/P24462/entry

[35] “UniProtKB P19099 · C11B2 HUMAN.” [Online]. Available: https://www.uniprot.org/uniprotkb/P19099/entry

[36] B. C. Chung, K. J. Matteson, R. Voutilainen, T. K. Mohandas, and W. L. Miller, “Human cholesterol side-chain cleavage enzyme, P450scc: cDNA cloning, assignment of the gene to chromosome 15, and expression in the placenta.” Proceedings of the National Academy of Sciences of the United States of America, vol. 83, no. 23, pp. 8962–8966, Dec. 1986. [Online]. Available: https://pmc.ncbi.nlm.nih.gov/articles/PMC387054/

[37] W. Tian, G. Wei, B. Duan, H. Peng, H. Liu, M. Zheng, Z. Lin, Z. Deng, Z. Zhang, and X. Qu, “Unveiling cytochrome P450 enzymes that catalyze steroid side-chain cleavage in bacteria,” Nature Communications, vol. 17, no. 1, p. 581, Dec. 2025. [Online]. Available: https://www.nature.com/articles/s41467-025-67278-3

[38] T. Padayachee, D. C. Lamb, D. R. Nelson, and K. Syed, “Structure–Function Analysis of the Self-Sufficient CYP102 Family Provides New Insights into Their Biochemistry,” International Journal of Molecular Sciences, vol. 26, no. 5, Feb. 2025. [Online]. Available: https://www.mdpi.com/1422-0067/26/5/2161

[39] A. Luthra, I. G. Denisov, and S. G. Sligar, “Spectroscopic features of cytochrome P450 reaction intermediates,” Archives of Biochemistry and Biophysics, vol. 507, no. 1, pp. 26–35, Mar. 2011. [Online]. Available: https://www.sciencedirect.com/science/article/pii/S0003986110005059

[40] F. P. Guengerich, “Roles of Individual Human Cytochrome P450 Enzymes in Drug Metabolism,” Pharmacological Reviews, vol. 76, no. 6, pp. 1104–1132, Oct. 2024.

[41] I. F. Sevrioukova and T. L. Poulos, “Understanding the mechanism of cytochrome P450 3A4: Recent advances and remaining problems,” Dalton Trans., vol. 42, no. 9, pp. 3116–3126, 2013. [Online]. Available: https://xlink.rsc.org/?DOI=C2DT31833D

[42] G. I. Lepesheva, L. Friggeri, and M. R. Waterman, “CYP51 as drug targets for fungi and protozoan parasites: Past, present and future,” Parasitology, vol. 145, no. 14, pp. 1820–1836, Dec. 2018.

[43] Y. Sun, R. Liu, Z. Luo, J. Zhang, Z. Gao, R. Liu, N. Liu, H. Zhang, K. Li, X. Wu, W. Yin, Q. Qin, X. Su, D. Zhao, and M. Cheng, “Identification of novel and potent triazoles targeting CYP51 for antifungal: Design, synthesis, and biological study,” European Journal of Medicinal Chemistry, vol. 280, p. 116942, Dec. 2024.

[44] R. Zhang, Y. Wang, A. Wu, J. Wang, and J. Zhang, “Strategies of targeting CYP51 for IFIs therapy: Emerging prospects, opportunities and challenges,” European Journal of Medicinal Chemistry, vol. 259, p. 115658, Nov. 2023.

[45] T. Padayachee, D. C. Lamb, D. R. Nelson, and K. Syed, “Structure-Function Analysis of the Biotechnologically Important Cytochrome P450 107 (CYP107) Enzyme Family,” Biomolecules, vol. 13, no. 12, p. 1733, Dec. 2023.

[46] S. Adam, L. Franz, M. Milhim, R. Bernhardt, O. V. Kalinina, and J. Koehnke, “Characterization of the Stereoselective P450 Enzyme BotCYP Enables the *In Vitro* Biosynthesis of the Bottromycin Core Scaffold,” Journal of the American Chemical Society, vol. 142, no. 49, pp. 20 560–20 565, Dec. 2020. [Online]. Available: https://pubs.acs.org/doi/10.1021/jacs.0c10361

[47] K. Suzuki, J. K. Stanfield, K. Omura, Y. Shisaka, S. Ariyasu, C. Kasai, Y. Aiba, H. Sugimoto, and O. Shoji, “A Compound I Mimic Reveals the Transient Active Species of a Cytochrome P450 Enzyme: Insight into the Stereoselectivity of P450-Catalysed Oxidations,” Angewandte Chemie - International Edition, vol. 62, no. 13, Mar. 2023.

[48] T. Hayashi, K. Harada, K. Sakurai, H. Shimada, and S. Hirota, “A role of the heme-7-propionate side chain in cytochrome P450cam as a gate for regulating the access of water molecules to the substratebinding site,” Journal of the American Chemical Society, vol. 131, no. 4, pp. 1398–1400, Feb. 2009.

[49] J. K. Stanfield, K. Omura, A. Matsumoto, C. Kasai, H. Sugimoto, Y. Shiro, Y. Watanabe, and O. Shoji, “Kristalle in Minutenschnelle: Sofortige Mikrokristallisation verschiedenster Varianten der CYP102A1-(P450BM3)-Hämdomäne,” Angewandte Chemie, vol. 132, no. 19, pp. 7681–7689, May 2020.

